# Implicit modeling of the conformational landscape and sequence allows scoring and generation of stable proteins

**DOI:** 10.1101/2024.12.20.629706

**Authors:** Yehlin Cho, Justas Dauparas, Kotaro Tsuboyama, Gabriel Rocklin, Sergey Ovchinnikov

## Abstract

Generative protein modeling provides advanced tools for designing diverse protein sequences and structures. However, accurately modeling the conformational landscape and designing sequences—ensuring that the designed sequence folds into the target structure as its most stable structure—remains a critical challenge. In this study, we present a systematic analysis of jointly optimizing *P*(structure|sequence) and *P*(sequence|structure), which enables us to find optimal solutions for modeling the conformational landscape. We support this approach with experimental evidence that joint optimization is superior for (1) designing stable proteins using a joint model (TrROS (TrRosetta) and TrMRF) (2) achieving high accuracy in stability prediction when jointly modeling (half-masked ESMFold pLDDT+ ESM2 Pseudo-likelihood). We further investigate features of sequences generated from the joint model and find that they exhibit higher frequencies of hydrophilic interactions, which may help maintain both secondary structure registry and pairing.

## Introduction

Generative modeling is becoming increasingly important in protein design, enabling the creation of diverse de novo structures with sequences that can refold into desired conformations [1–4]. When designing proteins, stability must be a top priority, as it enhances the efficiency of protein expression, purification, and crystallization [5]. Stability is primarily determined by a balance of forces that govern the free energies between the folded and unfolded states of the protein. It results from the intricate interplay between atomic interactions in the folded state and the higher conformational entropy in the unfolded state. Unstable proteins are prone to degradation and aggregation, making them unsuitable for biological or therapeutic applications. In an unfolded state, proteins can lose their function and potentially cause diseases [6, 7].

One primary approach to designing a folded protein is to generate a sequence that can fold into a specified target structure using structure-to-sequence models. These models (*P*(sequence|structure)) are often called “Inverse Folding” models, as they reverse the sequence-to-structure prediction process. However, many models labeled as such do not achieve true inverse folding. True inverse folding must satisfy two conditions: (i) the sequence must fold into the designed structure as its most energetically favorable configuration, and (ii) the sequence must not fold into any alternative structure with the same free energy [8]. A model trained with a *P*(sequence|structure) objective cannot see alternative conformations besides the one given, so there is a chance that the design sequences that fold into lower energy conformations other than the target structure (Figure 1A, left). Structure prediction models (*P*(structure|sequence)) can often be viewed as models that can see the entire energy landscape for a given sequence (Figure 1A, middle). However, most structure prediction models are trained on valid sequences known to fold, so the model inherently assumes that the sequences are good (i.e., has a high *P*(sequence)). As a result, the model may still regard a sequence as a good sequence, even if it is adversarial and does not fold well. For example, *P*(structure|sequence) models often struggle to differentiate between stable and unstable protein designs [9] and frequently fail when applied to protein design tasks alone [10, 11].

**Figure 1.**
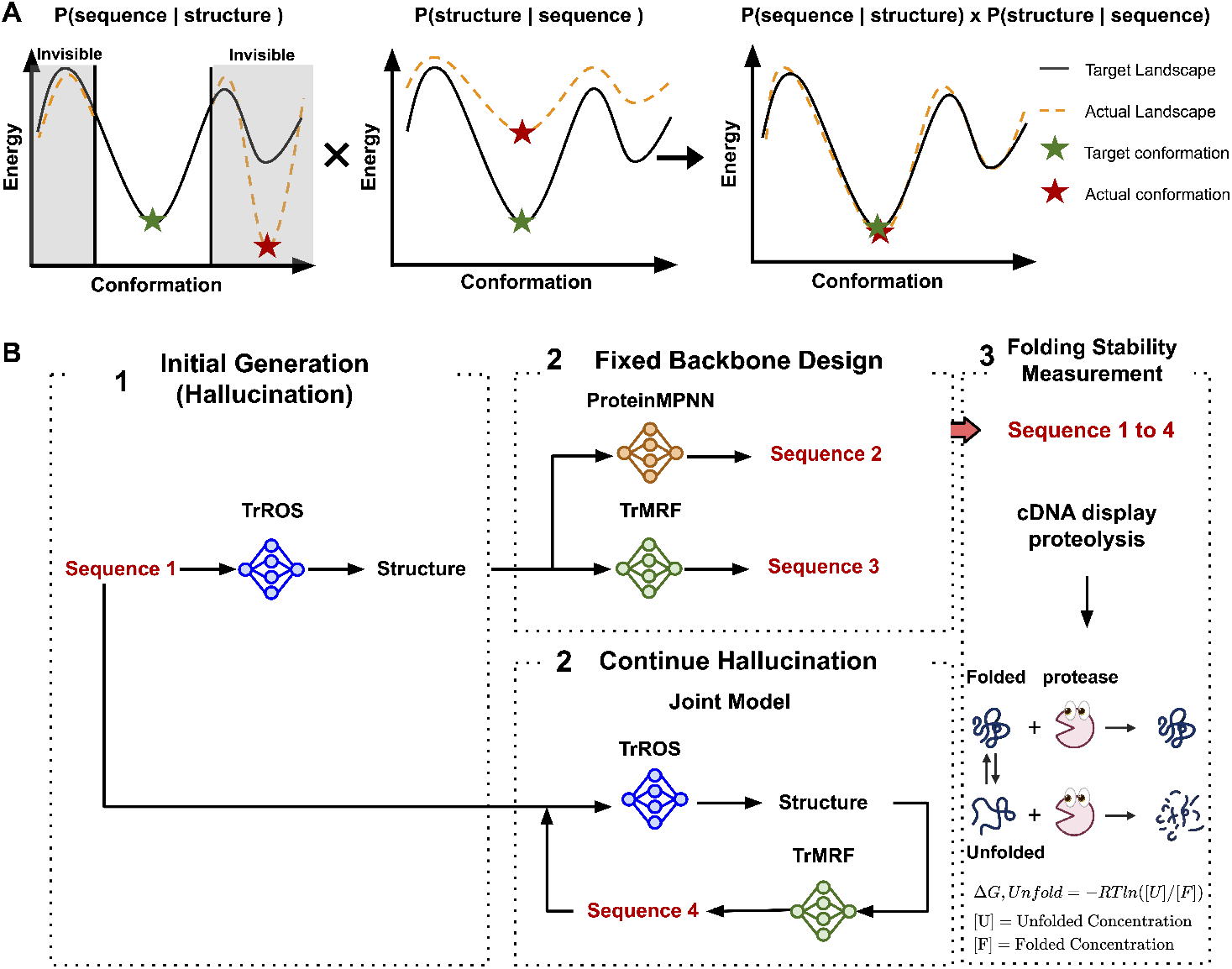
Overview of joint optimization in sequence and structure models: (A) (left) A sequence model predicts a sequence based on a given structure, targeting a specific conformation. (middle) A structure model predicts the structure from a given sequence. (right) A joint model integrates both sequence and structure information. (B) End-to-end sequence generation is performed using four different methods—TrROS, TrMRF, ProteinMPNN, and the Joint model (TrROS + TrMRF). After generating the sequences, protein stability was measured using the cDNA display proteolysis method.

Based on these observations, to achieve true “Inverse folding” we need to jointly optimize models that predict both *P*(sequence|structure) and *P*(structure|sequence), allowing the model to explore the full energy landscape. This approach optimizes the sequence towards the target structure while ensuring the target structure is the global minimum energy state (Figure 1 A, right). Here we provide a systematic analysis of (1) *P*(sequence|structure) (2) *P*(structure|sequence), and (3) the joint optimization of *P*(sequence|structure) and *P*(structure|sequence) using a four-model protocol, where each model takes one of (1)-(3) as its objective. We hypothesize joint optimization could perform better at modeling the conformational landscape and generating stable proteins, as it allows the backbone to be adjusted to accommodate the best possible sequence.

The four models are designed to generate four sequences that share the same fold but differ in sequence, optimizing based on different objectives. We measured the folding stability of designed protein sets using a cDNA display-based proteolysis method [12] to compare the stability of sequences designed from different objectives and to support our hypothesis. This generated folding stability dataset provides a unique opportunity to evaluate how well current models with varying objectives understand the protein landscape and stability. We also hypothesize that jointly optimizing both sequence and structure models will perform better at predicting and ranking stable proteins than using a single objective model. We approximate the confidence and likelihood scores of existing large protein models as a zero-shot predictor of stability. Our work underscores the importance of joint optimization of sequence and structure in modeling the conformational landscape to generate and score stable proteins.

## Results

### Joint optimization of sequence and structure models the conformational landscape and designs stable proteins

We investigate four models to generate de novo protein structures and sequences: TrROS, TrMRF, joint (TrROS + TrMRF), and ProteinMPNN (Figure 1 B). TrROS [13] is a model that generates the protein structure given a sequence (*p*(*structure*|*sequence*)). For unconditional protein generation to create thousands of structures and sequences, we use TrROS, starting with random amino acids and continuously applying gradient backpropagation until a clear distance matrix is obtained [13, 14]. The generated structures are then provided as input to ProteinMPNN and TrMRF, while the generated sequences are input to the joint model for generating new sequences (details are in Supplementary Figure SI 1). Cysteine is excluded to prevent the formation of disulfide bonds.

As a result, for each structure, we generated four sequences from four different models. Initially, we obtained a total of 20,668 sequences, with 5,167 sequences generated by each model. After filtering the sequences based on low AlphaFold2 inter-PAE (inter-chain predicted alignment error) to exclude proteins with potential aggregation or homo-oligomer formation, we retained 13,442 sequences, of which 5,708 have sequences from all four models. The structures generated by TrROS are diverse, with an average TM score of 0.4 between runs. Sequences with the same structures show low similarity across different models. The joint model and ProteinMPNN have the lowest average sequence similarity at 0.25, while TrROS and TrMRF have the highest at 0.407. To confirm whether sequences from the joint model, which continue hallucination from the output sequence of TrROS, could potentially alter the fixed backbone, we compared the differences in structure between sequences from the joint model and those from other models using RMSD (root mean square deviation). There are no significant differences in structure between the sequences of the joint model and those from other models, compared to the differences among the other models (Supplementary Figure SI 2).

We performed experimental validation by measuring protein folding stability using a high-throughput cDNA display proteolysis-based method [15–17]. We calculated the folding stability (D*G*) using the value *K*_50_, which is the protease concentration at which the cleavage rate is half the maximum rate, *k*_max_. *K*_50,*u*_ and *K*_50, *f*_ denote the unfolded and folded *K*_50_ values, respectively. *K*_50, *f*_ is experimentally measured from cDNA display experiments, while *K*_50,*u*_ is predicted by the NUTS model [12]. Further details can be found in the Supplementary Information.

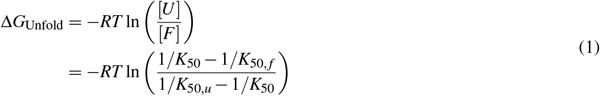

As shown in Figure 2A, the sequences generated by the joint model have the highest D*G*_,Unfold_ (which has the opposite sign of D*G*_,Fold_), representing the highest stability according to their highest resistance to two proteases, trypsin and chymotrypsin. To analyze the effect of sequence on folding stability, we compared sequences from different models that share nearly identical structures and calculated the difference in folding stability between the joint model sequences and those from other models. Sequences generated by the joint model (Figure 2B) exhibit higher folding stability than those from TrROS, TrMRF, and ProteinMPNN, with percentages of 80.5%, 74.4%, and 84.7%, respectively. This indicates that sequences designed by the joint model are more stable, even when structural components are disregarded.

**Figure 2.**
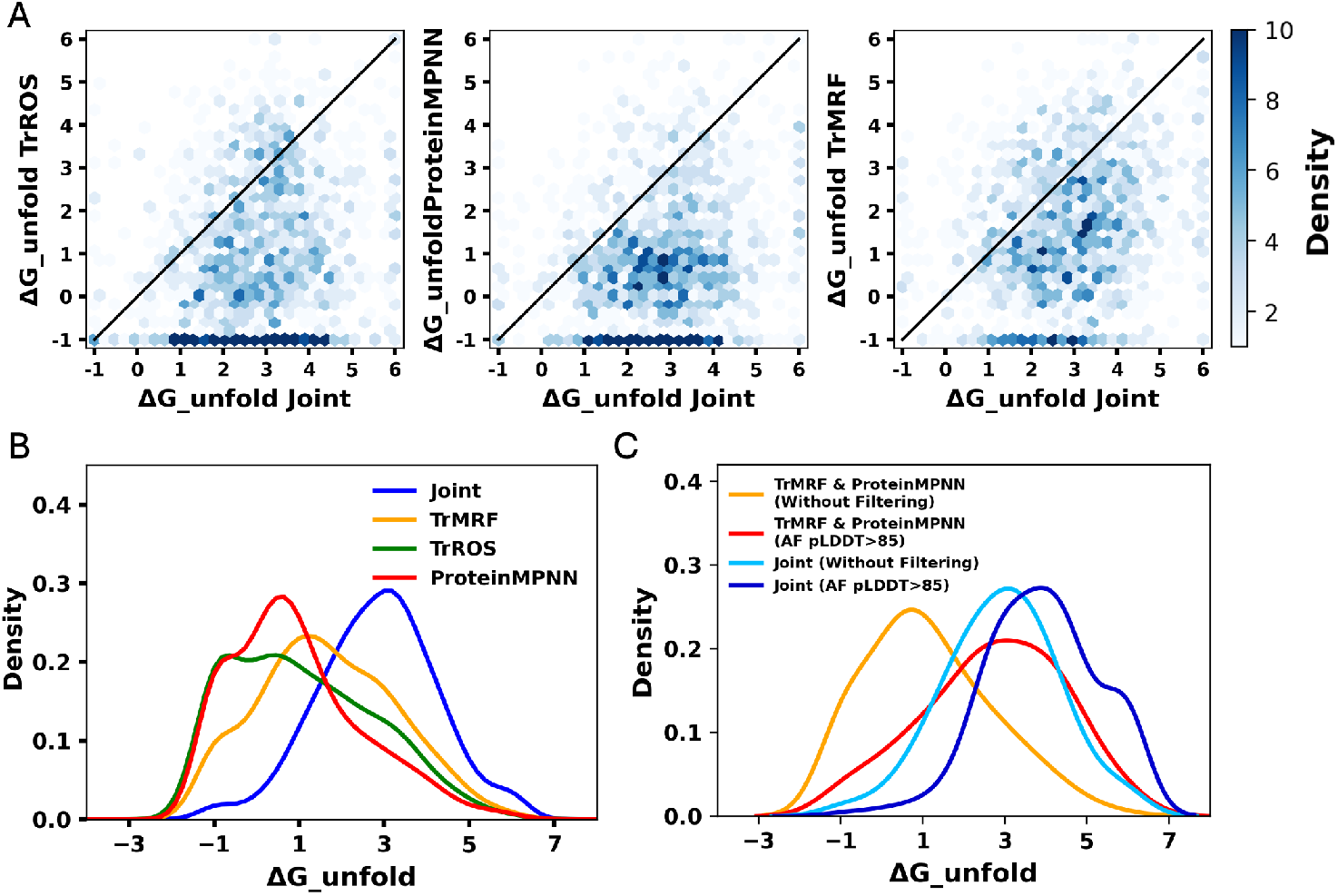
cDNA display proteolysis-based folding stability measurements show that the joint model generates more stable designs. (A) Comparison of D*G*_unfold_, shown in a hexaplot with the joint model on the x-axis and the single model on the y-axis, comparing the D*G*_unfold_ of sequences from each model that share the same structure. (B) Distribution of D*G*_unfold_ of four models. (C) Distributions of D*G*_unfold_ for sequences generated by (1) the joint model and (2) TrMRF and ProteinMPNN, with and without filtering for AF2 pLDDT *>* 85.

A common approach in protein design involves using a structure-to-sequence model to generate protein sequences and filtering these designed sequences based on computational confidence metrics, such as the AlphaFold2 (AF2) pLDDT score, which confirms that the designed proteins are well-folded with high confidence. For sequences generated from structure-to-sequence models (TrMRF and ProteinMPNN), we applied a filter of AF2 pLDDT *>* 85, where the AF2 prediction is based on the default setting of single-sequence prediction with three recycling steps. As shown in Figure 2C, the filtered sequences from the TrMRF and ProteinMPNN models exhibit a folding stability distribution similar to that of the joint model sequences without filtering. This is because, theoretically, designing a sequence based on a structure and then confirming its conformation using a structure prediction model can achieve the same objectives as the joint model, optimizing both *P*(structure|sequence) and *P*(sequence|structure). However, filtering joint model sequences with AF2 pLDDT *>* 85 shows an additional shift toward higher stability. This highlights that the joint model, by continuously adapting to both structural and sequential changes, may work better than any single-objective model by incorporating multiple objectives and designing highly stable designs when filtered with commonly used computational standards.

### Large sequence and structure models score folding stability in a zero-shot setting

Although filtering sequences based on common computational standards shifts the distribution toward more stable sequences compared to unfiltered ones, highly stable designs are not always achieved through common computational filtering (Figure 3A). Among all sequences with high folding stability (D*G >* 5), only 21.7% of sequences passed the AF2 (default setting, 3 recycles) and ProteinMPNN filters. A significant number of sequences exhibit moderate confidence values but high folding stability. Many highly stable designs still fall outside the filtering criteria, and given the heterogeneity of our dataset, we have the opportunity to explore alternative computational scoring matrices that may better correlate with folding stability. We assess how well current protein models rank stable versus unstable designs.

**Figure 3.**
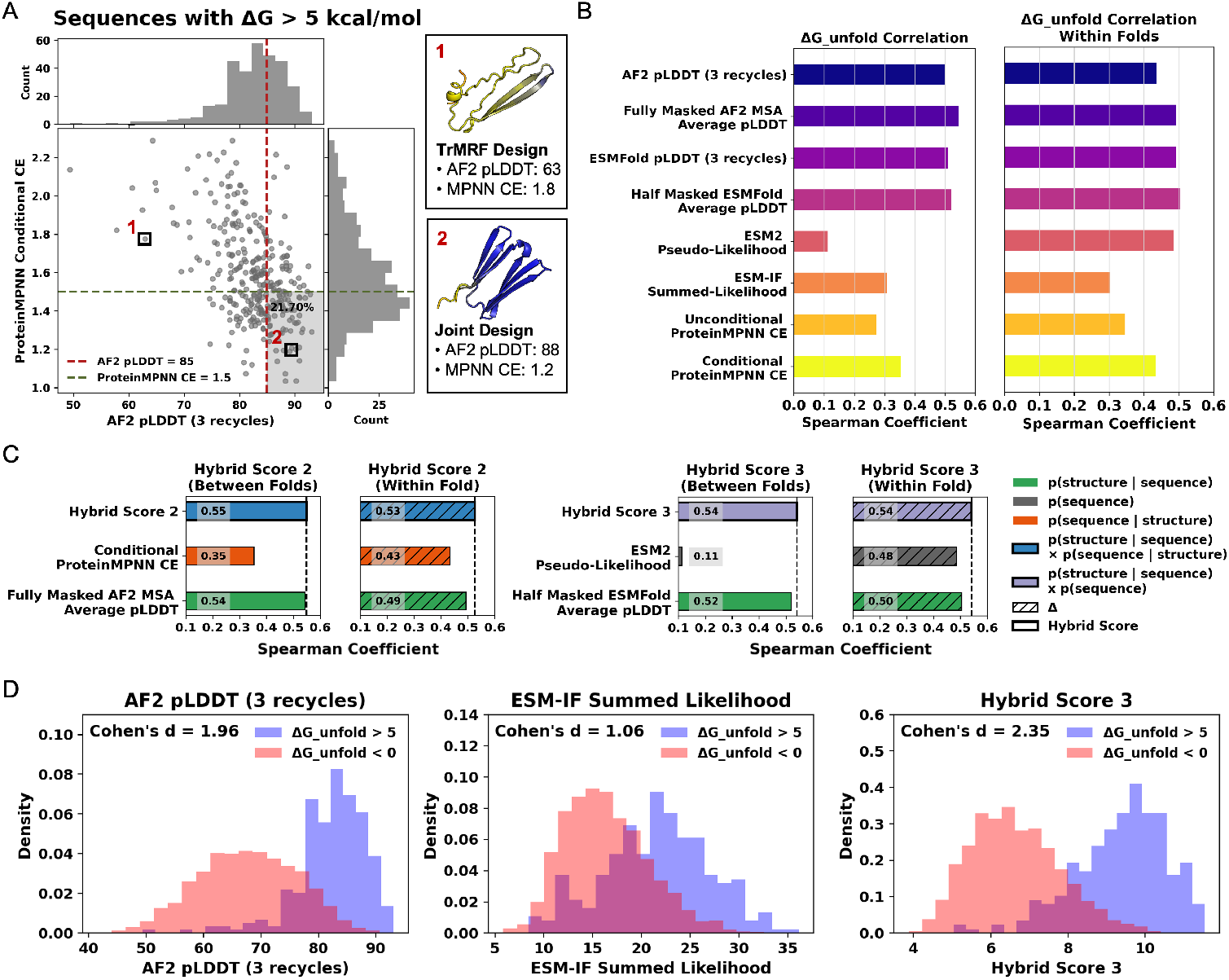
Zero-shot evaluation of large protein models identifies that hybrid scores combining sequence and structure-Based models achieve the best correlation with actual folding stability. (A) All sequences with D*G*_unfold_ *>*5 kcal/mol plotted with AF2 pLDDT on the x-axis and ProteinMPNN conditional CE on the y-axis. The gray box in the bottom right highlights sequences that passed common filtering standards: AF2 pLDDT *>* 85 and ProteinMPNN conditional CE *<* 1.5. (Upper) Example of a protein with low confidence but high folding stability, and (Lower) an example of a protein designed with high confidence and high stability. The structures are colored by per-residue level AF2 pLDDT value, where blue indicates high pLDDT and red indicates low pLDDT. (B) Spearman correlations between (Left) computational folding stability metrics and absolute folding stability(D*G*_unfold_) across different folds, and (Right) Spearman correlations between differences in absolute folding stability within folds and differences in metrics. (C) Spearman correlations between computational folding stability metrics and DG for both single and hybrid scores, along with correlations: (1) between folds and (2) differences within folds sharing the same structures. (D) Distribution of stable (D*G*_unfold_ *>*5 kcal/mol) and unstable designs (D*G*_unfold_ *<*0 kcal/mol) based on computational scoring metrics: AF2 pLDDT (3 recycles), ESM-IF Summed Likelihood, and Hybrid score 3.

Large protein models trained without fitness labels—such as activity, stability, or expression—have demonstrated the ability to predict and rank fitness metrics [18]. This capability may arise from the diverse, naturally occurring protein training datasets relevant to biological functions [19]. Zero-shot prediction enables the evaluation of a model’s ability to predict new tasks and classes without additional training [20, 21]. Similarly, we obtain model confidence and likelihood scores in a zero-shot setting and correlate them with actual stability scores to assess model performance. For sequence-conditioned models, we use the protein language model ESM-2 [22], ESMFold [23], and AF2. For structure-conditioned models, we use ESM-IF [24, 25] and ProteinMPNN [26]. We evaluate their ability to predict protein folding stability by calculating the Spearman correlation coefficient and analyzing the correlation between computational scores and experimental protein folding stability.

As shown in Figure 3B, AF2 and ESMFold pLDDT achieve the highest correlation with experimental absolute folding stability D*G*_Unfold_, as low pLDDT values are typically observed in unfolded segments. Compared to AF2 predictions with the default setting of three recycling steps, predictions without recycling demonstrate an improved overall correlation, as shown in Supplementary Figure SI 6. Additionally, fully masking the MSA input and averaging eight predictions with random seeds improves the Spearman correlation from 0.50 to 0.54 compared to providing the full sequence to the AF2 MSA module. ESMFold also achieves a higher correlation than the single-sequence approach by averaging eight pLDDT predictions with 50% random masking of the input sequence, which can enhance local stability predictions [27]. We reason that for de novo designs, language model features are less important, as they primarily capture evolutionary information. Therefore, masking language model features may improve AF2 and ESMFold predictions. While recycling enables the model to refine prediction quality, it can also result in unstable or poor-quality sequences being assigned high confidence.

Based on Figure 3B (left), it is clear that the ESM-2 model fails to predict the absolute folding stability compared to other models. We hypothesize that this limitation arises from the absence of a structural component in the model capable of evaluating structural stability alongside sequence stability. Instead of directly predicting folding stability, we conduct experiments to predict differences in protein folding stability across sequences derived from the same structure but generated using different models. Regarding the D*G*_Unfold_ correlation within folds, the ESM-2 pseudo-likelihood score shows a high correlation with the delta matrices (Figure 3B, right), in contrast to the low correlation observed between different folds. By comparing the stability of sequences within the same fold, we can offset the absolute differences between the two structures and assess the stability differences arising from sequence variations.

### Combining Sequence and Structure Models Improves Folding Stability Prediction

We introduce three hybrid scoring methods to enhance the accuracy of folding stability predictions by combining two of the following types of scores: (1) *p*(sequence) using ESM-2 Pseudo likelihood; (2) *p*(sequence | structure) using ProteinMPNN conditional cross-entropy (CCE); and (3) *p*(structure | sequence) using either AF2 or ESMFold pLDDT.

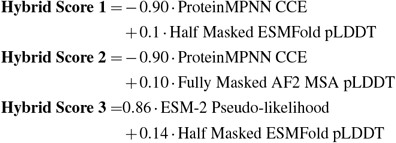

To determine the ratio between the two scores, we used 20% of the datasets to find the ratio that achieves the best Spearman correlation with experimental absolute stability and the hybrid score, and we obtained the final Spearman correlation tested on the remaining 80% of the dataset. Hybrid Score 2, which combines the fully masked AF2 MSA average pLDDT and ProteinMPNN CCE, and Hybrid Score 3, which combines the pseudo-likelihood of ESM-2 with a 50% masked ESMFold average pLDDT, achieve the best correlation with experimental folding stability scores (Figure 3C). For Hybrid Score 3, optimizing both *p*(*structure*|*sequence*) and *p*(*sequence*), allows for adjusting the *p*(*structure*|*sequence*) model, which intrinsically assumes *p*(*sequence*) = 1, meaning all sequences are equally likely. By incorporating *p*(*sequence*), we can more accurately assess the likelihood of both the sequence and its corresponding structure. Compared to the commonly used AF2 filtering (3 recycles) and the ESM-IF summed likelihood score—presented as a zero-shot absolute protein folding stability predictor [25]—the Hybrid score more effectively distinguishes stable and unstable designs, as shown in Figure 3D. This supports our hypothesis that combining structure- and sequence-based scores can improve the generation and prediction of stable proteins.

### Joint Model Designs Sequences with Higher Frequency of Hydrophilic Interactions

To understand the differences between sequences generated by the joint model and single-objective models, we focus on amino acid pairs in contact, as interactions between amino acids are crucial for maintaining both secondary and global structure. We identified these amino acid pairs by measuring the distances between heavy atoms in side chains that are less than 5 Å and separated by more than five residues in the predicted structures from AlphaFold2. As a control, we evaluated the contact pairs in 200 PDB native structures. As shown in Figure 4A, sequences from the TrROS, TrMRF, and joint models exhibit enriched interactions of charged, polar, and hydrophobic residues, with same-charge pairs showing negative couplings [12, 28]. Notably, sequences from the joint model show a higher log odds ratio for both hydrophilic residue pairs (Figure 4A). As shown in Figure 4B, the secondary structure of the joint model sequence is better maintained compared to those from other single models, forming electrostatic and hydrogen bonds between side chains. Charged residues, such as lysine and glutamic acid, are sampled more frequently in joint models than in TrMRF models (Figure 4C), with a high probability of lysine-glutamic acid pairs forming contacts (Figure 4D), along with other hydrophilic interactions. These features may contribute to the high stability of joint model sequences, allowing them to maintain both their secondary and global structures.

**Figure 4.**
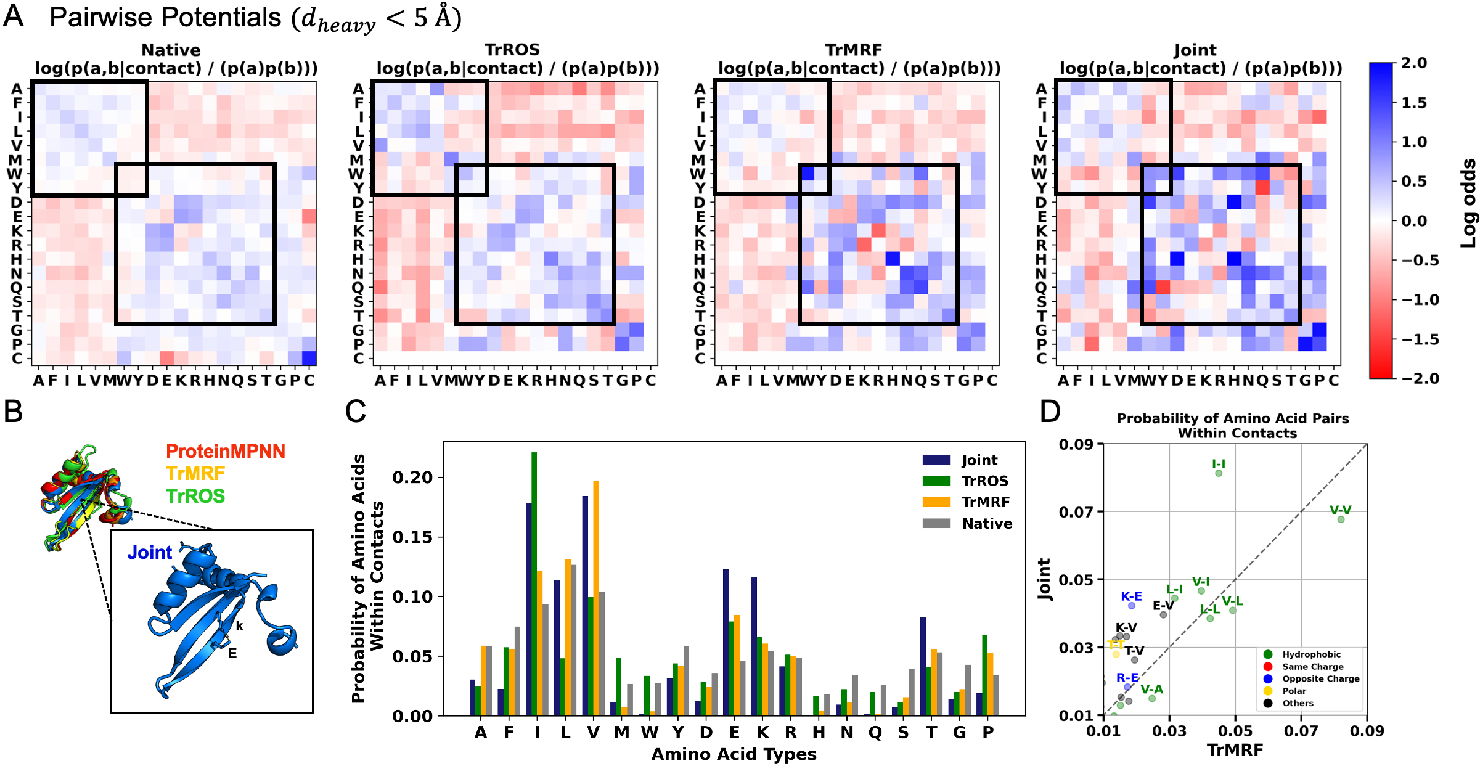
Sitewise and pairwise analyses show that the joint model samples a higher frequency of hydrophilic interaction pairs. (A) 20×20 amino acid pairwise potentials in contacts where distances between heavy atoms are less than 5 Å for joint model-sampled sequences. The pairwise potential is calculated as log(P(a, b | contact)) / P(a)P(b). Black boxes indicate interactions between charged and polar residues and hydrophobic residues. Red denotes enriched interactions, while blue represents depleted interactions. For the native, pairwise potentials were calculated from 200 PDB native sequences. (B) AlphaFold2-predicted structures from sequences generated by the joint, TrROS, TrMRF, and ProteinMPNN models, with a zoomed in view showing electrostatic and hydrogen bonds formed between side chains in the joint model structure. (C) Bar plots show the probability of single amino acids within contacts from the joint model, TrROS, TrMRF, and native proteins. Cysteine is excluded from the plots, as it is not used in the sequence design. (D) Scatter plots showing the probability of amino acid pairs within contacts between the Joint model and TrROS model sequences.

## Discussion

Our systematic analysis of (1) *P*(structure|sequence), (2) *P*(sequence|structure), and (3) joint optimization of both demonstrates that joint optimization implicitly models the conformational landscape, generating the most stable proteins by simultaneously designing the sequence and adjusting the backbone to fit the sequence. Joint model sequences exhibit a folding stability distribution similar to those generated by *P*(sequence|structure) models filtered with the common AF2 pLDDT standard, which models *P*(structure|sequence). This similarity arises because filtering *P*(sequence|structure) sequences with *P*(structure|sequence) helps exclude sequences that fold into alternative states by globally exploring the conformational landscape, performing a sequential form of joint optimization. However, after applying the same AF2 pLDDT filter, joint model sequences exhibit greater stability compared to those from *P*(sequence|structure) models, suggesting that the joint model simultaneously optimizes both sequence and structure to find an optimal solution.

We also evaluate the ability of the current models to score stability in a zero-shot setting. We compared the correlation between AF2 and ESMFold stability predictions across multiple different settings and found that, for de novo proteins, masking language model information—by masking the AF2 MSA input, masking the ESMFold input sequence, and averaging multiple outputs—helps increase the correlation with experimental stability. For AF2, decreasing recycling also prevents the model from having high confidence in adversarial sequences. Hybrid scores, derived from combining sequence and structure-based models, exhibit the highest correlation with experimental folding stability compared to any single model. Notably, the hybrid score combining (1) AF2 MSA average pLDDT and ProteinMPNN CCE, with (2) ESM2 pseudo-likelihood and half-masked ESMFold pLDDT, achieves the best results. By considering both (1) *P*(sequence|structure) and *P*(structure|sequence), and (2) *P*(sequence) and *P*(structure|sequence), we can assess the likelihood of both the sequence and its corresponding structure, thereby enhancing the hybrid score for predicting stability.

In conclusion, joint optimization enables implicit modeling of the conformational landscape and sequence design, facilitating the scoring and generation of stable proteins. Our absolute stability dataset offers a unique opportunity to evaluate different models and analyze stability across sequences sharing the same folds, allowing for the independent analysis of sequence components from the structure. Although our current work primarily evaluates mini-proteins of fewer than 80 amino acids, which may limit its applicability to larger, multi-domain proteins, we expect that this methodological analysis and unique dataset will lay the foundation for designing stable proteins and for optimizing and evaluating models in conformational landscape modeling.

## Methods

### TrROS, TrMRF, and Joint Models

TrROS integrates sequence information to estimate conservation and coevolution features calculated from the inverse covariance matrix of the input sequences. These features are then used as inputs to predict protein structures using ResNet blocks. ResNet blocks, featuring dilated convolutions and dropout layers, enable the model to capture long-range dependencies within protein sequences. Finally, the prediction head outputs 6D features, including torsion angles, distance predictions, backbone torsion angles, and omega angles. Rosetta then reconstructs the 3D structure of the protein using 6D features. For unconditional protein generation, we initialize the model with a random amino acid sequence, predict the structure, and backpropagate the gradients through the TrRosetta model. Based on the KL (Kullback–Leibler divergence) loss between the predicted and the background structure distance map (averaged over all protein structures), we can optimize both the amino acid sequences and the structures simultaneously. This process continues until a structure with a clear distance matrix is obtained.

TrMRF is an inverted model of TrROS; more specifically, for a given structure, it predicts the conservation and coevolution features and maximizes the pseudo-likelihood of the associated Multiple Sequence Alignment (MSA), approximating the probability of the sequence given the structure (*p*(*sequence*|*structure*)). ResNet layers with varying dilation rates enable the model to capture features at different spatial resolutions. After the ResNet blocks, the model branches into a separate path to extract pairwise potentials and long-range dependencies. This path reshapes the output of the ResNet into a higher-dimensional tensor representing coevolution and conservation features. During sequence sampling with the TrMRF model, sequence optimization begins with a random amino acid sequence, and the categorical cross-entropy is computed between this sequence and the predicted sequences from the TrMRF model. The loss updates the random amino acid sequence, but it is not backpropagated into the conservation or coevolution matrices or the predicted sequence. The sampling process continues until the loss converges.

The joint model combines the TrROS and TrMRF models. Without retraining the model weights, we utilize the pre-trained TrROS and TrMRF models. For sequence generation, we jointly optimize the loss from both the TrROS and TrMRF models and update the sequence accordingly. Unlike the TrMRF and ProteinMPNN models, which perform fixed backbone design by starting from the structure provided by the TrROS model, the joint model begins with the final sequence of the predicted structure. Then it uses this sequence as the initial input to predict 6D features using the TrROS model. These 6D features are given as input to the TrMRF model, which generates sequences. There are two losses in the optimization process: TrMRF loss and TrROS loss. TrMRF loss is backpropagated into the TrMRF model and the input sequence, while TrROS loss is backpropagated through the TrROS model. The optimization process continues until the loss no longer decreases.

### Data Filtration

For generated sequences, we observe that some designs have hydrophobic patches that can form homo-oligomers. Therefore, even if one protein is proteolyzed, the other can bind to the fragmented protein, leading to a false signal. To computationally distinguish proteins that can potentially form homo-oligomers, we use the predicted alignment error (inter-PAE) between chains provided by AlphaFold2 [29, 30]. We provide two copies of proteins to AlphaFold2 and let it predict whether they form homo-oligomers using the inter-PAE value. Typically, inter-PAE *>* 15 indicates that the protein exists as a monomer [29].

### Sequence Based Zero-Shot Method for Predicting Protein Stability

**ESM 2** is a protein language model that enables the direct prediction of masked positions of various protein attributes, including structure and function, from an individual sequence[22]. ESM-2 is trained on the UniRef database [31] by predicting 15% of the masked amino acids in input protein sequences. The pseudo-likelihood of ESM-2 approximates the probability of a sequence *P*(*sequence*), and serves as a metric to assess the performance of a sequence model in representing data [22, 32, 33]. Since the ESM attention matrix and representations can predict contacts within the protein [34, 35], we assume that pseudo-likelihood might help distinguish between stable and unstable structures. We calculate the ESM-2 pseudo-likelihood for our generated sequence *S* of length *L* by masking each position *s*_*i*_ with the [MASK] token, denoted as *S*_*\i*_ := (*s*_1_,…, *s*_*i-*1_, [*MASK*], *s*_*i*+1_,…, *s*_*L*_). This calculation is performed by aggregating the conditional log probabilities log *P*_*ESM*2_(*s*_*i*_|*S*_*\i*_) for each token.

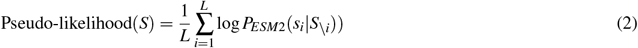

**ESMFold** utilizes the ESM-2 language model and structure module to produce precise structure predictions for proteins [23]. ESMFold achieves accurate predictions of the structures of de novo proteins using a single sequence. It demonstrates the capability to differentiate proteins into groups that are designable and those that are un-designable[27]. ESMFold processes a protein sequence (*S*) of length (*L*), generating the predicted local distance difference test (*pLDDT*) and *distogram logits* for information about contact predictions. pLDDT is a matrix that assesses the quality of a protein structure at a local level, while distogram logits (*g*(*S*)) represent the structural probability distribution *p*(*xyz*|*S*). We hypothesize that calculating the difference between predicted and actual protein structures using cross-entropy, which measures the disparity between the predicted structure distribution from the model and the true structure, could help classify protein stability by evaluating the confidence in the predicted structure. Cross-entropy, *H*(*F, g*(*S*)), is calculated by digitizing the actual structure into an *L* × *L* × *M* matrix, where *M* is the number of bins.

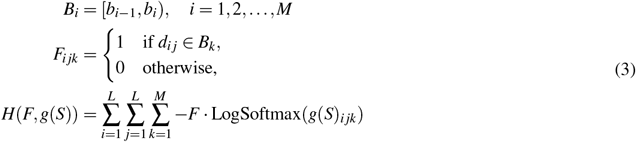

The number of bins, denoted as *M*, is determined by dividing the total range, which is the range of pairwise contact distances, *b*_*M*_ *b*_1_, by the width of the bin, *h*. This can be expressed as 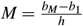. Each bin, *B*_*i*_, represents an interval [*b*_*i-*1_, *b*_*i*_). The matrix *F* encodes whether the value of the distance *d*_*ij*_ between the *i*th and *j*th positions of *C*_*β*_ belongs to a bin *B*_*i*_. It assigns a one if it does and a 0 otherwise.

### Structure Based Zero-Shot Method for Predicting Protein Stability

**ESM-IF** Previous works showed that the ESM-IF model can be used to predict the absolute folding stability of proteins [24, 25]. To evaluate folding stability, the authors used the output logits from the ESM-IF decoder to assess the likelihood of amino acids at each position. This likelihood is determined by applying the Softmax function to the 20 amino acid logits at each position and then summing across the length of the protein to obtain the overall folding stability.

**ProteinMPNN** generates a sequence from protein backbone coordinates with a message-passing structure consisting of an encoder-decoder framework [26, 36]. ProteinMPNN incorporates protein backbone coordinates by representing protein residues as nodes and establishing nearest neighbor edges by considering pairwise distances between backbone atoms. These input features are subsequently passed through three encoder layers and then into the decoder layers to obtain a sequence. ProteinMPNN predicts the probability of a sequence (logits) given a structure that is in the opposite direction compared to ESMFold. We employ the ProteinMPNN cross-entropy term, *H*(*X, f* (*X*)), as a criterion for assessing the folding stability. This involves calculating the cross-entropy between the ground-truth sequence and the logits. This choice is based on the observation that ProteinMPNN can evaluate the quality of ROSETTA decoy datasets of 133 native protein structures. The cross-entropy of the protein decoy aligns well with the TM score between the decoy and the ground truth structure, as illustrated in.

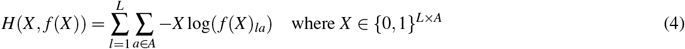

*X* is a one-hot encoded protein sequence of length *L* and composed of *A* different types of amino acids. Logits from the model can be generated either by conditioning on the ground truth sequence during decoding, resulting in conditional logits, or without conditioning on any sequence, termed unconditional logits. We term the cross-entropy derived from conditional logits as *conditional cross-entropy* (CCE) and that from unconditional logits as *unconditional cross-entropy* (UCE).

## Acknowledgements

We thank the members of the Rocklin and Ovchinnikov labs for useful discussions. Y.C. acknowledges funding from SBS scholarships and Takeda Fellowship. S.O. acknowledges funding from NIH grant DP5OD026389, NSF MCB2032259, and Amgen.

## References

1. Wu, Z., Johnston, K. E., Arnold, F. H. & Yang, K. K. Protein sequence design with deep generative models. Curr. opinion chemical biology 65, 18–27 (2021).

2. Ingraham, J. B. et al. Illuminating protein space with a programmable generative model. Nature 1–9 (2023).

3. Jumper, J. et al. Highly accurate protein structure prediction with alphafold. Nature 596, 583–589 (2021).

4. Watson, J. L. et al. De novo design of protein structure and function with rfdiffusion. Nature 620, 1089–1100 (2023).

5. Deller, M. C., Kong, L. & Rupp, B. Protein stability: a crystallographer’s perspective. Acta Crystallogr. Sect. F: Struct. Biol. Commun. 72, 72–95 (2016).

6. Stein, A., Fowler, D. M., Hartmann-Petersen, R. & Lindorff-Larsen, K. Biophysical and mechanistic models for disease-causing protein variants. Trends biochemical sciences 44, 575–588 (2019).

7. Yue, P., Li, Z. & Moult, J. Loss of protein structure stability as a major causative factor in monogenic disease. J. molecular biology 353, 459–473 (2005).

8. Yue, K. & Dill, K. A. Inverse protein folding problem: designing polymer sequences. Proc. Natl. Acad. Sci. 89, 4163–4167 (1992).

9. Kim, T.-E. et al. Dissecting the stability determinants of a challenging de novo protein fold using massively parallel design and experimentation. Proc. Natl. Acad. Sci. 119, e2122676119 (2022).

10. Goverde, C. A., Wolf, B., Khakzad, H., Rosset, S. & Correia, B. E. De novo protein design by inversion of the alphafold structure prediction network. Protein Sci. 32, e4653 (2023).

11. Verkuil, R. et al. Language models generalize beyond natural proteins. bioRxiv 2022–12 (2022).

12. Tsuboyama, K. et al. Mega-scale experimental analysis of protein folding stability in biology and design. Nature 620, 434–444 (2023).

13. Norn, C. et al. Protein sequence design by conformational landscape optimization. Proc. Natl. Acad. Sci. 118, e2017228118 (2021).

14. Anishchenko, I. et al. De novo protein design by deep network hallucination. Nature 600, 547–552 (2021).

15. Park, C., Zhou, S., Gilmore, J. & Marqusee, S. Energetics-based protein profiling on a proteomic scale: identification of proteins resistant to proteolysis. J. molecular biology 368, 1426–1437 (2007).

16. Park, C. & Marqusee, S. Pulse proteolysis: a simple method for quantitative determination of protein stability and ligand binding. Nat. methods 2, 207–212 (2005).

17. Yamaguchi, J. et al. cdna display: a novel screening method for functional disulfide-rich peptides by solid-phase synthesis and stabilization of mrna–protein fusions. Nucleic acids research 37, e108–e108 (2009).

18. Notin, P. et al. Proteingym: Large-scale benchmarks for protein fitness prediction and design. Adv. Neural Inf. Process. Syst. 36 (2024).

19. Roney, J. P. & Ovchinnikov, S. State-of-the-art estimation of protein model accuracy using alphafold. Phys. Rev. Lett. 129, 238101 (2022).

20. Lampert, C. H., Nickisch, H. & Harmeling, S. Learning to detect unseen object classes by between-class attribute transfer. In 2009 IEEE conference on computer vision and pattern recognition, 951–958 (IEEE, 2009).

21. Radford, A. et al. Language models are unsupervised multitask learners. OpenAI blog 1, 9 (2019).

22. Lin, Z. et al. Language models of protein sequences at the scale of evolution enable accurate structure prediction. BioRxiv 2022, 500902 (2022).

23. Lin, Z. et al. Evolutionary-scale prediction of atomic-level protein structure with a language model. Science 379, 1123–1130 (2023).

24. Hsu, C. et al. Learning inverse folding from millions of predicted structures. In International conference on machine learning, 8946–8970 (PMLR, 2022).

25. Cagiada, M., Ovchinnikov, S. & Lindorff-Larsen, K. Predicting absolute protein folding stability using generative models. bioRxiv 2024–03 (2024).

26. Dauparas, J. et al. Robust deep learning–based protein sequence design using proteinmpnn. Science 378, 49–56 (2022).

27. Hermosilla, A. M., Berner, C., Ovchinnikov, S. & Vorobieva, A. A. Validation of de novo designed water-soluble and transmembrane proteins by in silico folding and melting. bioRxiv 2023–06 (2023).

28. Anishchenko, I., Ovchinnikov, S., Kamisetty, H. & Baker, D. Origins of coevolution between residues distant in protein 3d structures. Proc. Natl. Acad. Sci. 114, 9122–9127 (2017).

29. Mirdita, M. et al. Colabfold: making protein folding accessible to all. Nat. methods 19, 679–682 (2022).

30. Akdel, M. et al. A structural biology community assessment of alphafold2 applications. Nat. Struct. & Mol. Biol. 29, 1056–1067 (2022).

31. Suzek, B. E. et al. Uniref clusters: a comprehensive and scalable alternative for improving sequence similarity searches. Bioinformatics 31, 926–932 (2015).

32. Shin, J., Lee, Y. & Jung, K. Effective sentence scoring method using bert for speech recognition. In Asian Conference on Machine Learning, 1081–1093 (PMLR, 2019).

33. Salazar, J., Liang, D., Nguyen, T. Q. & Kirchhoff, K. Masked language model scoring. arXiv preprint 1910.14659 (2019).

34. Rao, R., Meier, J., Sercu, T., Ovchinnikov, S. & Rives, A. Transformer protein language models are unsupervised structure learners. Biorxiv 2020–12 (2020).

35. Fung, A., Koehl, A., Jagota, M. & Song, Y. S. The impact of protein dynamics on residue-residue coevolution and contact prediction. bioRxiv 2022–10 (2022).

36. Ingraham, J., Garg, V., Barzilay, R. & Jaakkola, T. Generative models for graph-based protein design. Adv. neural information processing systems 32 (2019).

37. Sosnick, T. R. The folding of single domain proteins-have we reached a consensus? Biophys. J. 100, 373a (2011).

38. Coey, C. T. & Drohat, A. C. Kinetic methods for studying dna glycosylases functioning in base excision repair. In Methods in enzymology, vol. 592, 357–376 (Elsevier, 2017).

