## Supplementary Information for "Implicit modeling of the conformational landscape and sequence allows scoring and generation of stable proteins"

### A. cDNA Display-based Folding Stability Measurement

#### 1. Analyzing protein folding stability via proteolysis in the cDNA display framework

In the case of many small, single-domain proteins, folding can be simplified as a two-state system. This system involves two distinct populations of proteins—those in a folded state and those in an unfolded state—separated by a free-energy barrier [37]. Folding stability, in this context, is measurable as the difference in free energy levels between the folded and unfolded states, denoted as the folding free energy ($\Delta G_{Fold}$). This can be experimentally measured using K50, where the unfolded state is denoted as $K50_U$ and the folded state as $K50_F$. Tsuboyama, Kotaro, et al. introduce a predictive tool for assessing the folding stability of unfolded proteins ($K50_U$) under protease conditions, called No U-Turn Sampler (NUTS), using a dataset comprising approximately 64,238 protein scramble proteolysis samples [12]. The model can distinguish stable sequences from unstable ones through protease treatment. Furthermore, a robust correlation emerges between ΔG values obtained from cDNA display proteolysis and known folding ΔG values [12]. To validate the folding stability of our designed proteins, we conduct a DNA display-based proteolysis experiment using two different proteases: trypsin, which selectively acts on basic amino acids, and chymotrypsin, which targets amino acids with aromatic properties.

#### 2. Experiment Details of high-throughput display-based cDNA proteolysis

To measure the folding stability of proteins, we used proteolysis reactions. We assumed single-turnover kinetics [38] with a substantial excess of enzyme compared to the substrate. Equation (5) describes a proteolysis process in which the protease (E) and the folded protein (F) can form an enzyme-substrate complex (FE), leading to proteolysis and degradation into a product (P). Furthermore, the enzyme can proteolyze the unfolding protein (U) to form an unfolded enzyme-substrate complex (UE), resulting in degradation and the production of a decorated protein (P). There is an equilibrium between the folded and unfolded states of the protein, and the free energy between the two states represents the folding stability ΔG in equation (6). The bi-directional arrows indicate the reversible process.

$$\begin{gathered} E + F \rightleftharpoons FE \rightarrow E + P \\ E + U \rightleftharpoons UE \rightarrow E + P \end{gathered} \tag{5}$$

$$\Delta G, Unfold = -RT \ln\left(\frac{[U]}{[F]}\right) \tag{6}$$

If $\Delta G_{,\text{Unfold}}$ is positive, it indicates that the folded state is more stable than the unfolded state. The calculation of $\Delta G_{,\text{Unfold}}$ relies on the concentrations of unfolded and folded proteins, specifically using the ratio $[U]/[F]$. This ratio can be obtained the measured sequence-specific K50 (the protease concentration at which the cleavage rate is half of the maximum rate, kmax), unfolded state (K50,U), and folded state (K50, F), as shown in equation (7). Here, we set K50, F to be same for all sequences, because we assume that cleavage in the folded state occurs exclusively in the constant regions.

$$\begin{aligned} K_{50} \approx K_d = \frac{[E][S]}{[ES]} &= \frac{([U]+[F])[E]}{([UE]+[FE])} \\ &\approx \frac{[U]+[F]}{[U]/K_{50,u} + [F]/K_{50,f}} \end{aligned} \tag{7}$$

$$\frac{[U]}{[F]} \approx \frac{1/K_{50} - 1/K_{50,f}}{1/K_{50,u} - 1/K_{50}} \tag{8}$$

Proteolysis reactions are conducted under two conditions: with trypsin and with chymotrypsin. For each group, experiments are performed across 11 different protease concentrations, arranged in a threefold dilution series, with one condition without protease to determine the point at which the cleavage rate is half of Kmax (K50)[12]. Bayesian inference is employed to derive K50 and ΔG values for all sequences within our library. This analysis comprises two main models. The first model, referred to as the 'K50 model,' calculates the K50 values for each sequence based on the count data from sequencing. The second model, named the 'unfolded state model', anticipates the unfolded state K50 value (K50,U) for each sequence by considering its actual sequence [12]. As shown in Figure SI 4, there is a clear linear correlation between ΔG values of trypsin and chymotrypsin.

### B. Details of Folding Stability Predicting Models

We plot one-to-one comparison scatter plots between individual prediction scores (Figure SI 7). The x-axis includes the half-masked ESMFold pLDDT, full and half-masked ESMFold Distogram cross-entropy, ESM2 pseudo-likelihood, AlphaFold2 pLDDT, ESM-IF summed likelihood, Unconditional ProteinMPNN CE, Conditional ProteinMPNN CE, and hybrid scores 1, 2, and 3. The y-axis is fixed to Full sequence ESMFold pLDDT.

### C. Spearman comparisons between TM (template modeling score) and CE (cross entropy) of ProteinMPNN

We hypothesize that ProteinMPNN, which is trained on structure space from the PDB dataset, can evaluate the difference between proteins in their lowest energy conformation and decoys created using relaxation or backrub methods. We can approximate how well the ground truth sequence matches the given structure by calculating the cross-entropy loss between the output logits and the ground truth sequence. To test this hypothesis, we prepared 134 targets. For each target, we sampled 1,000 Rosetta decoy structures [19]. We then provided these sampled structures into the ProteinMPNN model, predicted logits, and calculated the categorical cross-entropy loss between these logits and the wild-type sequence.

When we plotted the cross-entropy (CE) and TM scores between decoy and native structures, a clear inverse linear correlation emerged: as cross-entropy decreases, the similarity between the two structures increases. With a total of 134 targets and 1,300 decoy structures per target, we observed a high Pearson correlation, as shown in Figure SI 8 and Figure SI 9. Three representative structures showed relatively low correlations, between 0.4 and 0.5. These proteins either have disordered regions, causing sampled decoys from Rosetta to have a low TM score with the wild-type structure, or the disordered regions result in different conformations while sampling the decoys (Figure SI 10).

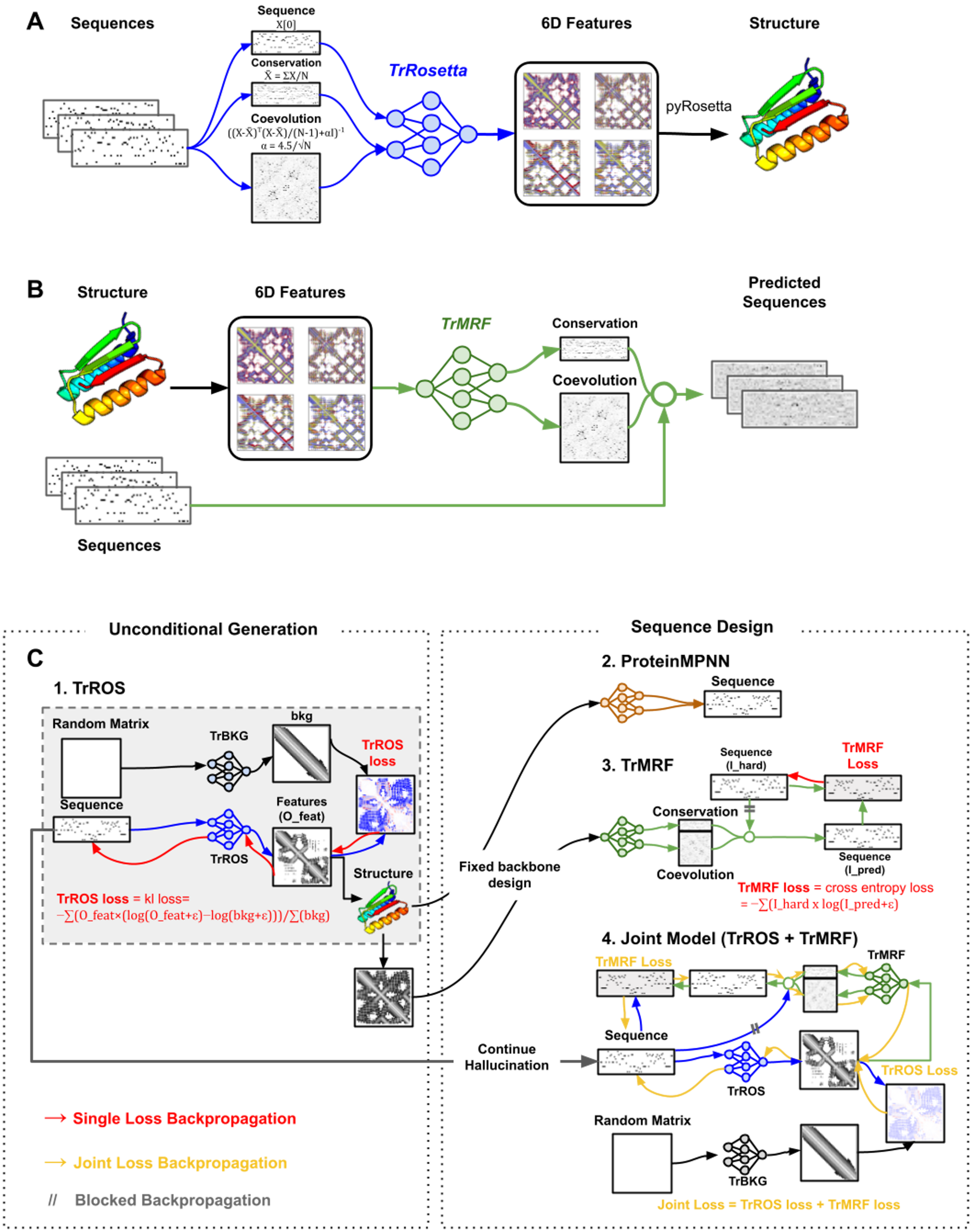


**Figure SI 1.** De novo Protein Sequence and Structure Generation: (A) Model architecture of TrROS, (B) Model architecture of TrMRF, and (C) Overview of end-to-end generation of sequences using four different models. $I_{\text{true}}$ and $I_{\text{pred}}$ are the sequence logits given and predicted, $O_{\text{feat}}$ represents the predicted 6D features, and bkg refers to the background 6D features.

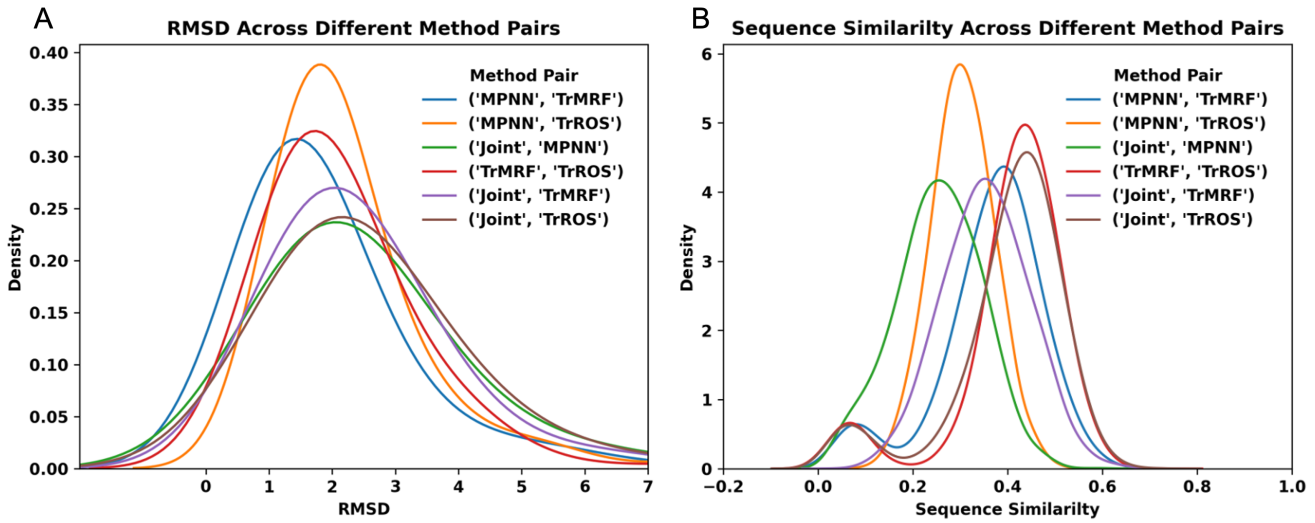


**Figure SI 2.** (A) Structure similarity measured by RMSD across different pairs of methods for structures filtered by AlphaFold pLDDT scores higher than 70 and inter-PAE higher than 15 (a total of 5,167 structures). (B) Sequence similarity across different pairs of methods (a total of 20,668 sequences)

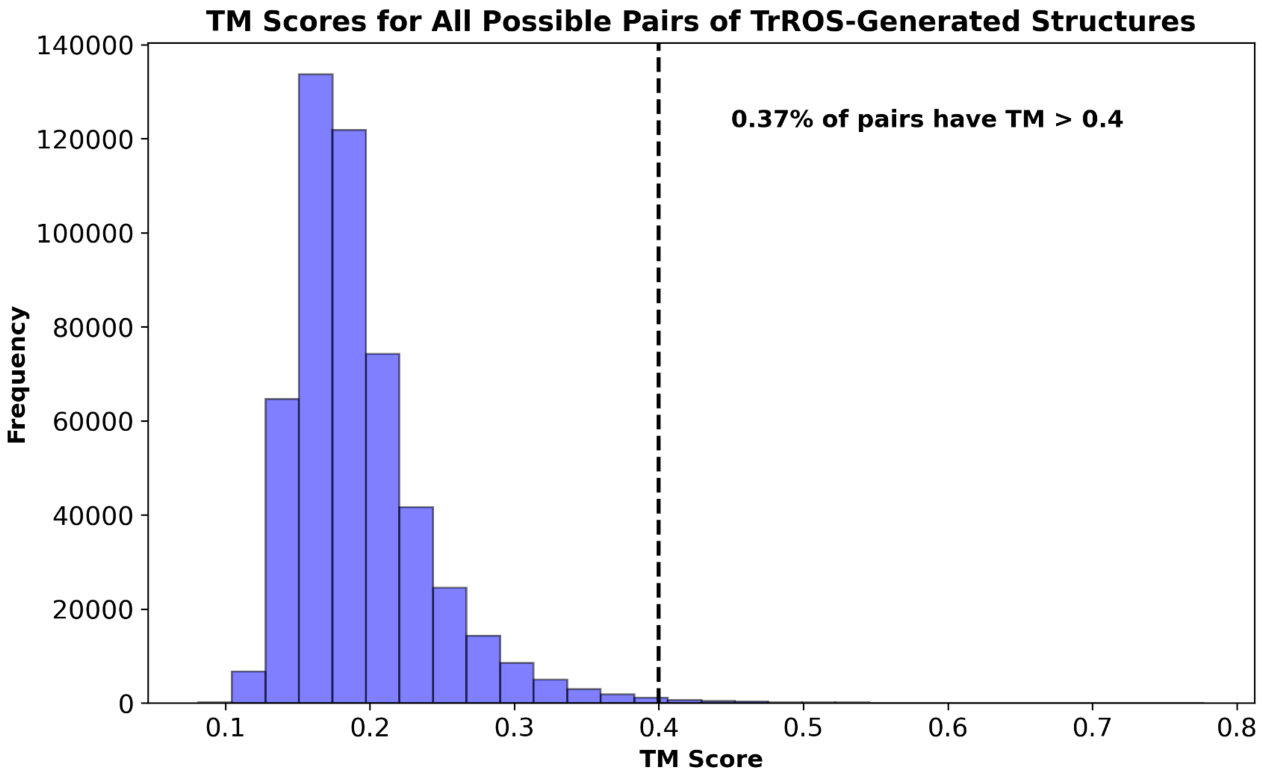


**Figure SI 3.** TM scores between TrROS-generated structures: Generally, if the TM score is less than 0.4, the two proteins are not considered to be in the same fold. Here, 0.37% of pairs have a TM score greater than 0.4, so most of them are different

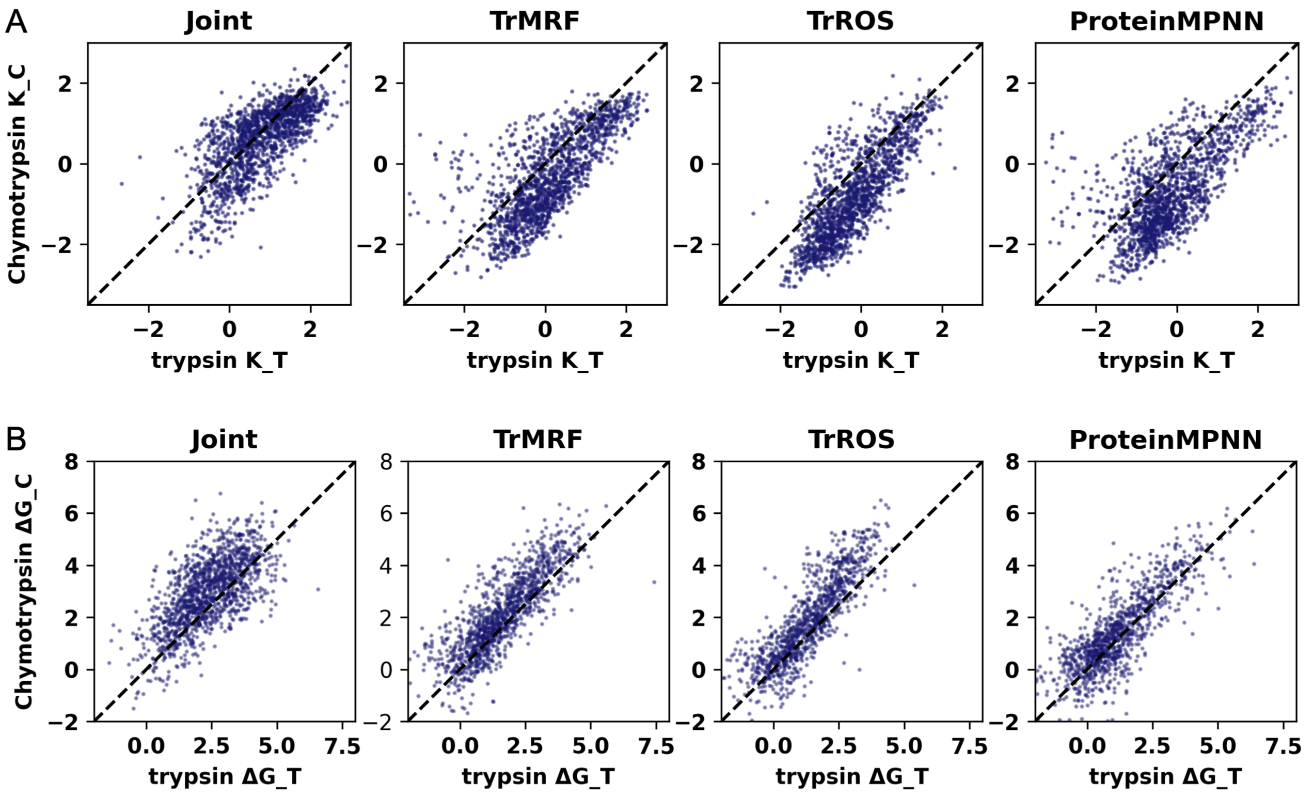


**Figure SI 4.** Comparisons of (A) resistance to trypsin ($k_T$) vs. chymotrypsin ($k_C$) and (B) folding stability to trypsin ($\Delta G_T$) vs. chymotrypsin ($\Delta G_C$).

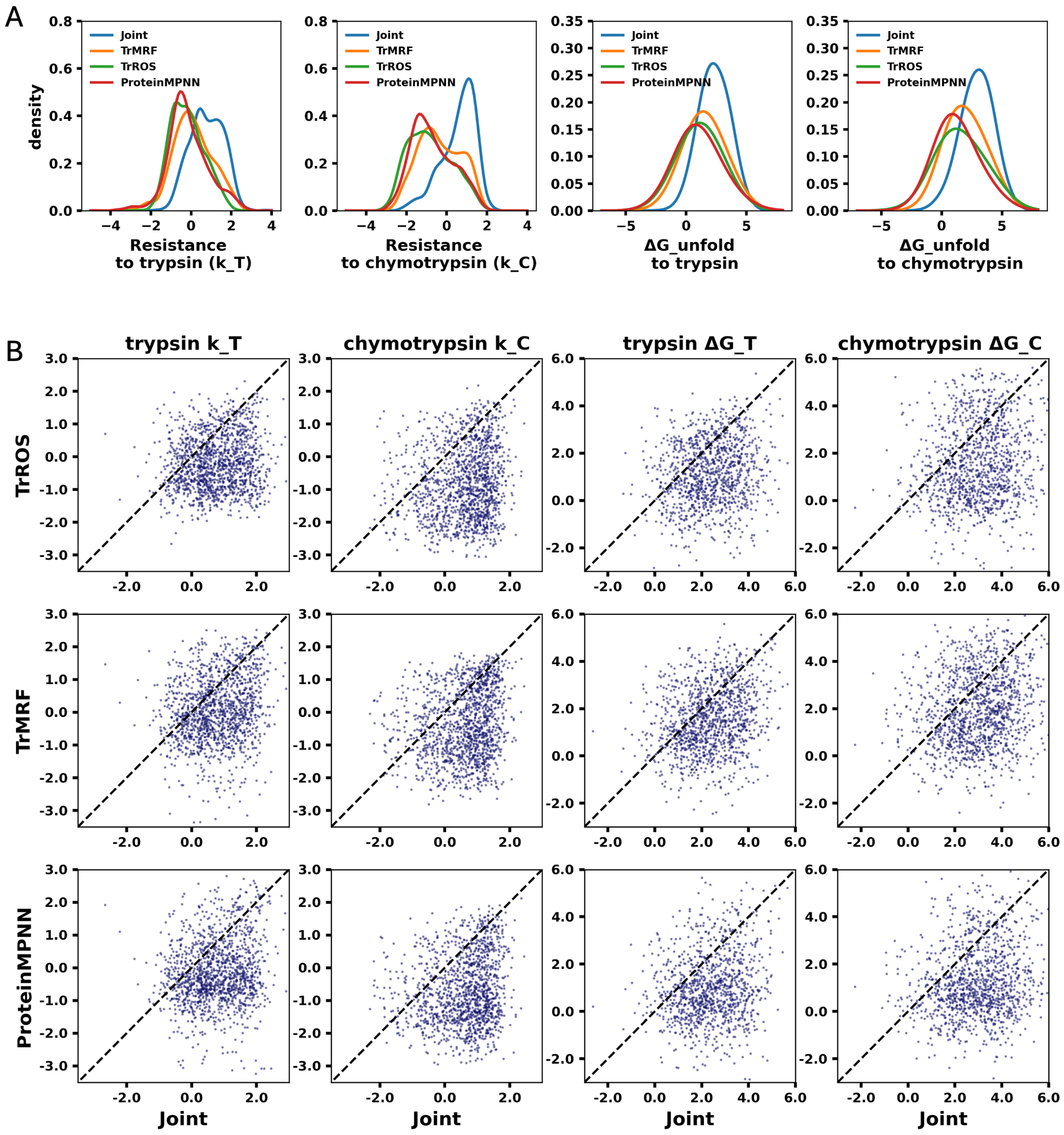


**Figure SI 5.** Experimental metrics comparisons of folding stability between different models. (A) resistance to trypsin ($k_T$), chymotrypsin ($k_C$), folding stability to trypsin ($\Delta G_T$), and chymotrypsin ($\Delta G_C$). (B) Folding stability comparisons between single models and the joint model0 using the same metrics as in (A).

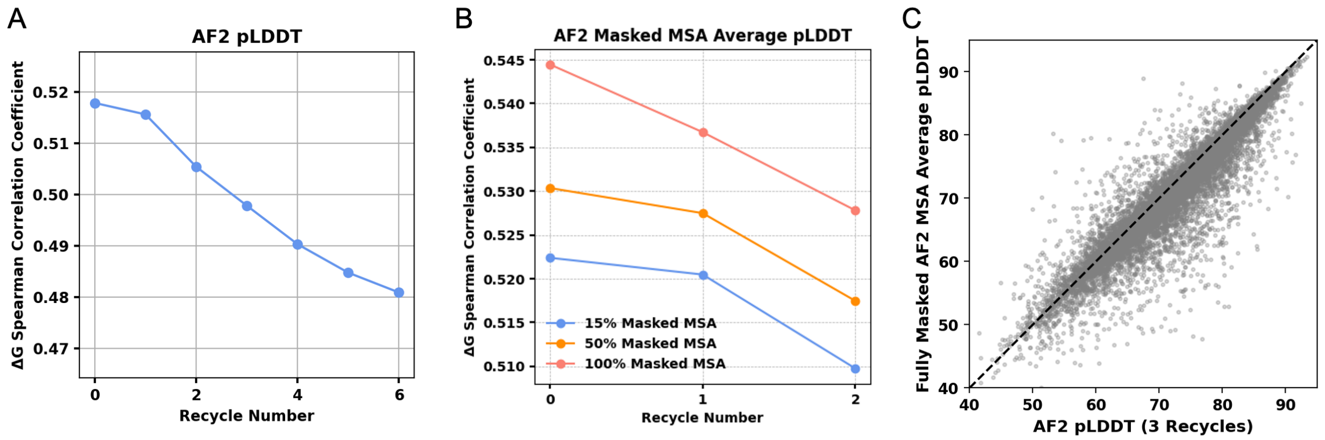


**Figure SI 6.** Correlation between AF2 pLDDT and experimental folding stability under different conditions: (A) varying recycle numbers, (B) varying percentages of MSA masking, and (C) comparison of AF2 pLDDT predictions with 3 recycles versus fully masked MSA input without recycling.

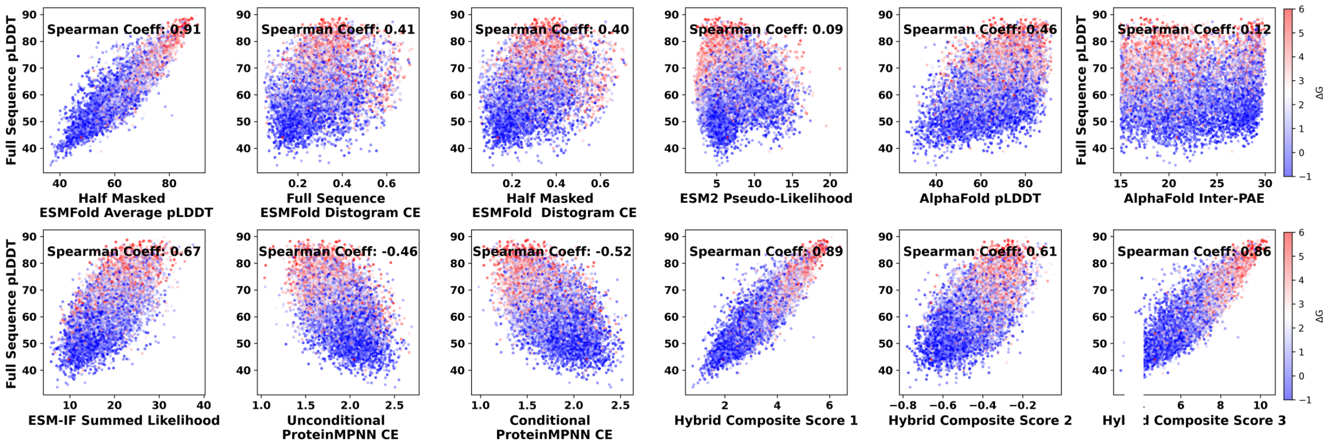


**Figure SI 7.** Comparison of the models' folding stability predictions in a zero-shot setting with the full-sequence-based ESMFold pLDDT score. The plotted data points are colored by $\Delta G_{\text{unfold}}$ values, with red indicating positive values and blue indicating negative values.

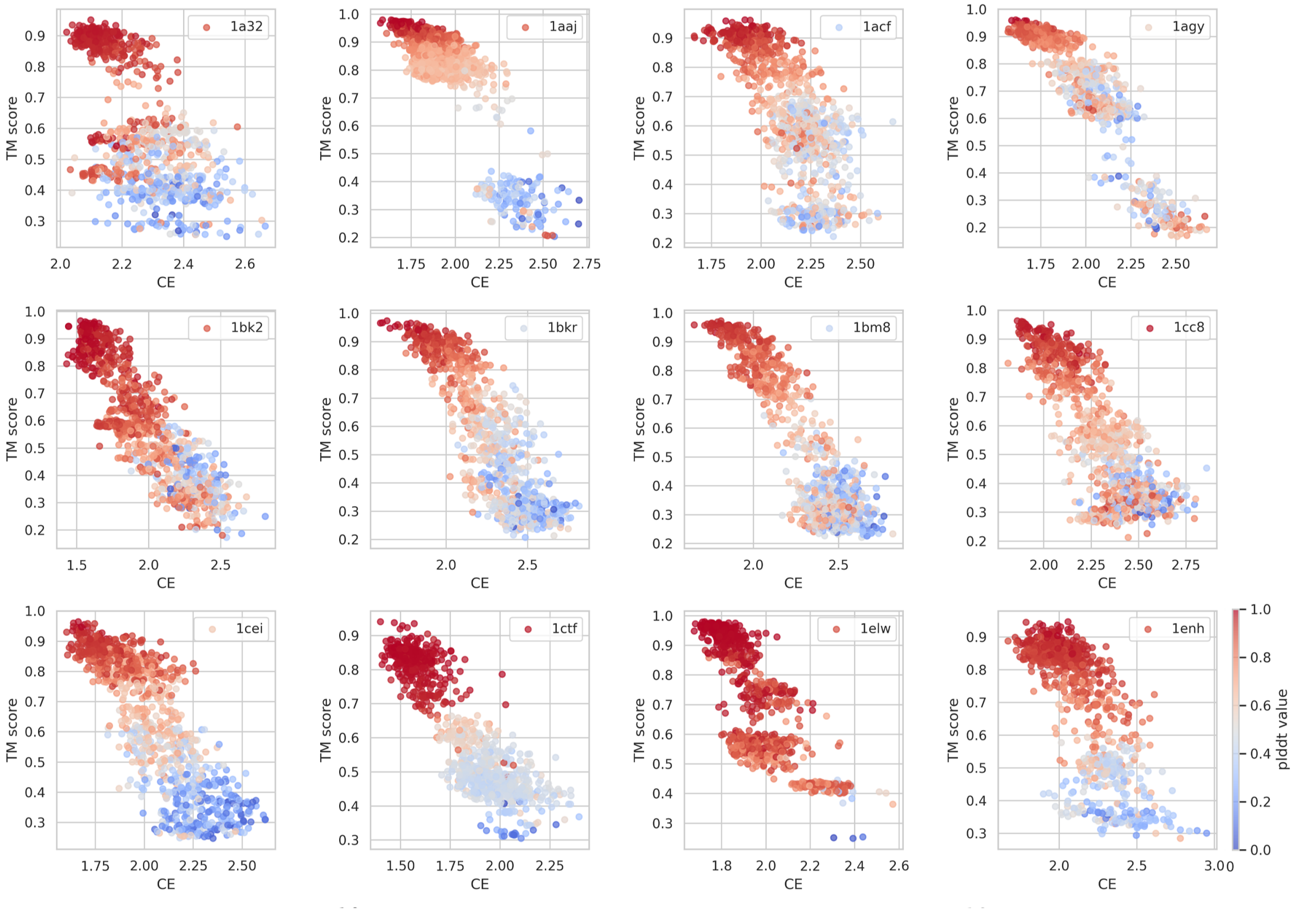


**Figure SI 8.** Scatter plot of TM score and ProteinMPNN cross-entropy (CE), colored by Alphafold2 pLDDT.

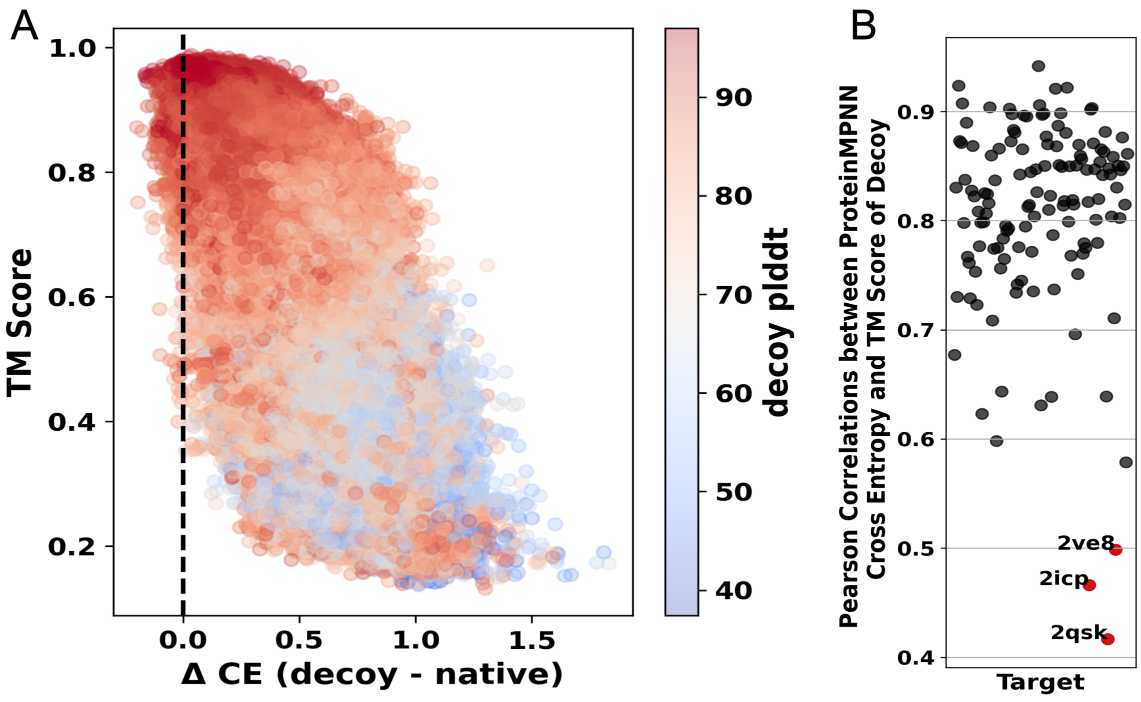


**Figure SI 9.** Correlation of TM score and CE for(A) total decoys (B) each individual target decoy.

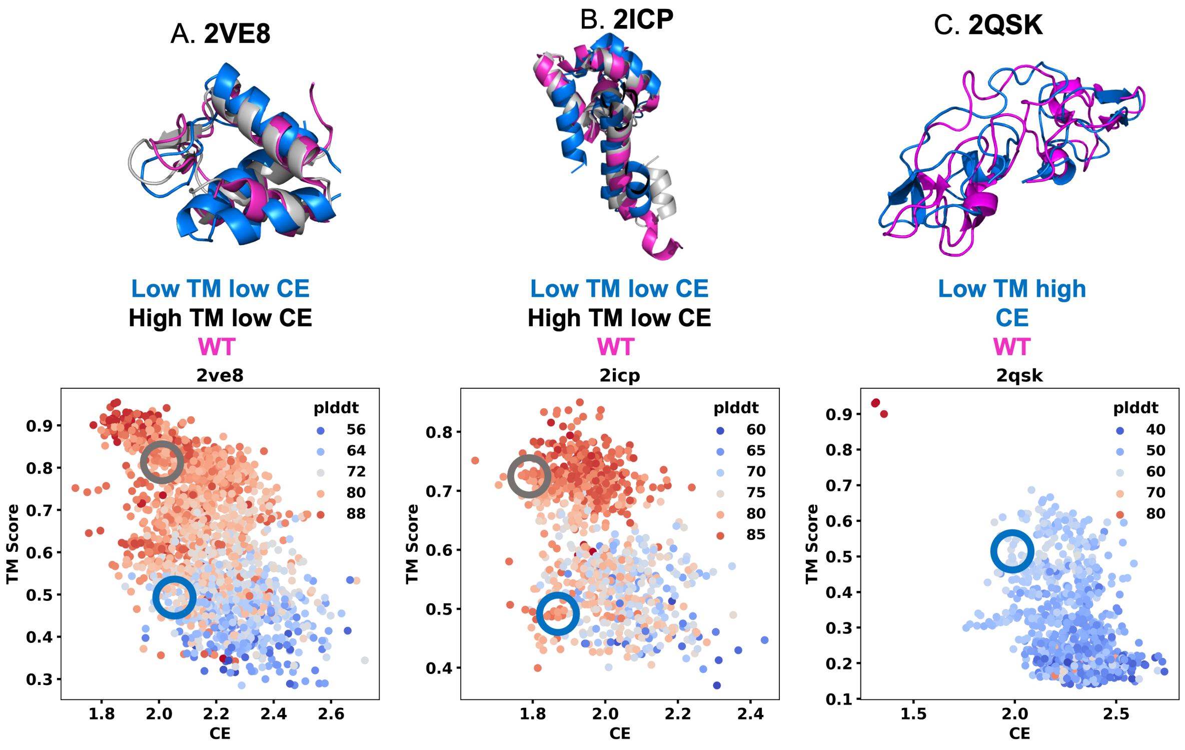


**Figure SI 10.** Three representative structures show less correlation between the TM score and ProteinPNN cross entropy (CE): A. 2VE8, B. 2ICP, C. 2QSK structures. The top figures show overlapped structures of the wild type and decoy structure, indicating a high TM score and a low TM score with the wild type.

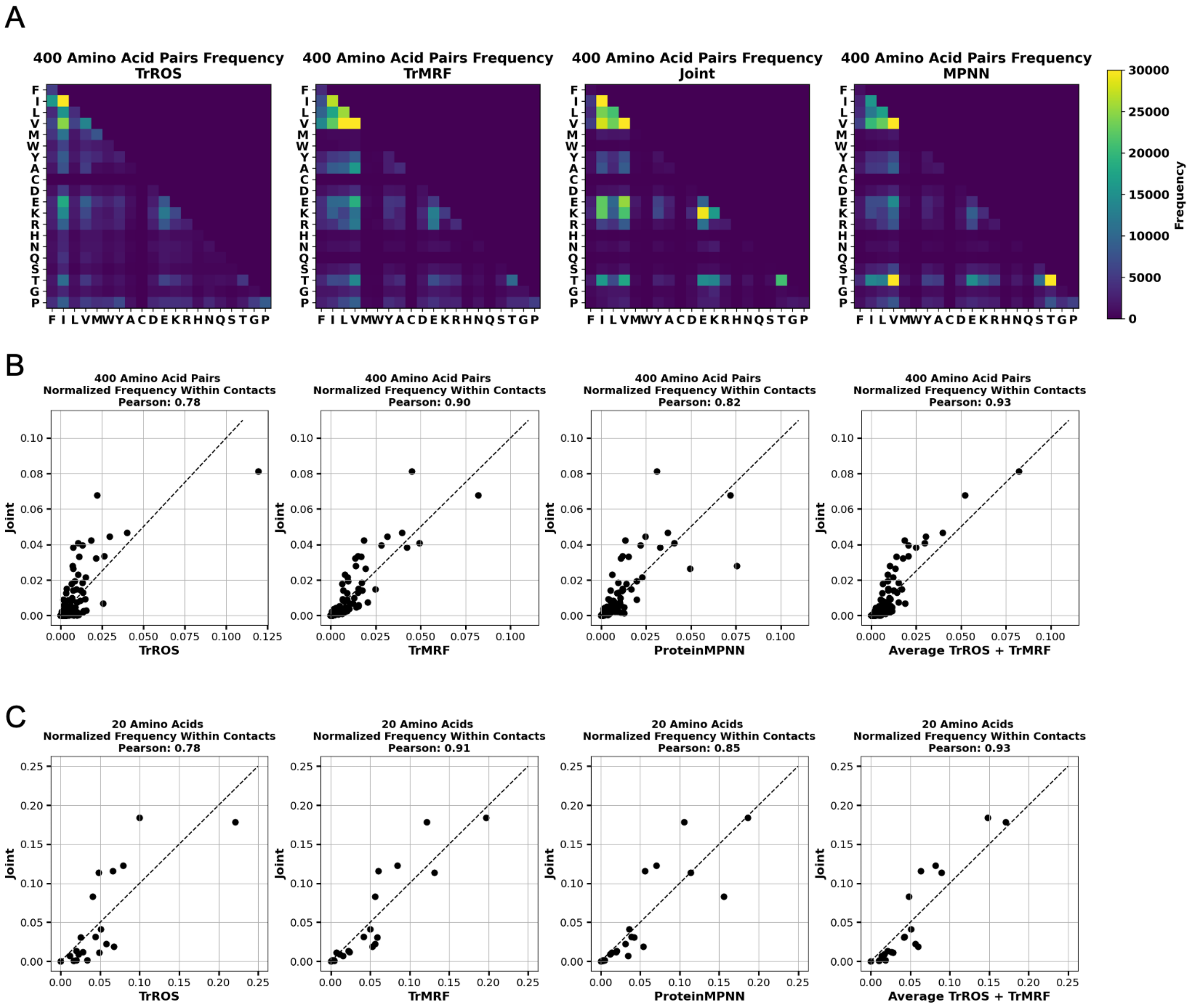


**Figure SI 11.** Amino Acid Frequency in Contacts. (A) Frequency of 20x20 amino acid pairs in contacts across sequences generated from four different models. Scatter plots compare (B) the normalized frequency of 400 amino acid pairs and (C) the frequency of 20 single amino acids between the single model and the joint model.
